# Identify the origin and end cells and infer the trajectory of cellular fate automatically

**DOI:** 10.1101/2022.09.28.510020

**Authors:** Zehua Zeng, Cencan Xing, Lei Hu, Shah Roshan, Yuanyan Xiong, Hongwu Du

## Abstract

Cellular state identification and trajectory inference enable reconstructions of cell fate dynamics from single-cell RNA sequencing. However, the identification of cell fate trajectories requires a large number of computational resources or known biological process, and lack a method to alleviate both of these deficiencies at the same time. Here, we present scLTNN, a method that automatically infers origin and end cell state from scRNA-seq data and calculates the developmental trajectory and differentiation direction of cells with only a few computational resources and time consummation. We apply scLTNN to disentangling subpopulation kinetics in CD8+ T cell, pancreatic endocrinogenesis, and the development of zebrafish embryos. scLTNN displays a strong trajectory inference ability cross-species. scLTNN features a modular design that can be flexibly extended to any scRNA-seq analysis task. The complete package is available online at https://github.com/Starlitnightly/scltnn.

## Introduction

Cell fate trajectories can help people understand various biological processes such as disease evolution, life development, and cell regulation^1–3^. In contrast, single-cell transcriptomics allows for cellular heterogeneity under transient slices of cell fate trajectories (including cellular differentiation and genealogical selection) at single-cell resolution^4^. The resulting computational problems are referred to as trajectory inference and cell state identification^5,6^. Cell fate trajectory inference algorithms aim to reorder cells from cell heterogeneity and construct a sequence of fate trajectories^7,8^. Based on the concept of trajectory inference, the algorithms are divided into two main aspects: the dynamics inference algorithm based on RNA velocity of spliced-to-unspliced mRNA ratio^4,9^, and the graph evolution algorithm based on the expression of key genes^10,11^. However, each of these two strategies has its deficiencies: the RNA velocity-based algorithm relies on a large number of computational resources as well as time consumption, and the key gene-based graph algorithm relies on the setting of manual a priori knowledge (artificial selection of developmental root and direction), while the identification of developmental root is often tricky in cancer as well as in cell subpopulations.

Here, we introduce scLTNN (scRNA-seq latent time neural network), a modular framework for the latent time of cell estimation, origin and end cell identification, and cell fate trajectory computation (Fig.1b). We combine the advantages of two strategies for trajectory inference: 1) using a pre-trained neural network model to analyze the single-cell expression matrix and estimate the latent time associated with RNA velocity, thus reducing computational resources and time consumption (Fig.1a,1c). 2) using genes whose expression patterns correlate with gene counts and the previously determined latent time to automatically filter cell origin and end states, thus avoiding the artificial selection of the starting point and direction of cell trajectories. scLTNN can achieve an efficient and accurate inference of cell fate trajectories. In addition, we constructed an LTNN time by combining the normal distribution characteristics of origin, middle and end cell times and the Dweibull distribution characteristics of diffusion pseudotime, whose distribution characteristics are similar to the latent time calculated by RNA velocity and have highly similar pseudotime results. We demonstrate LTNN’s capabilities on pancreatic lineage development in mice, CD8+ T cell development in humans, axial mesoderm lineage of the zebrafish embryo, and correctly identify the origin and end cell states and cell fate trajectory cross-species. scLTNN is available as a scalable, user-friendly, open-source software package with documentation and tutorials at https://github.com/Starlitnightly/scltnn.

**Figure 1.**
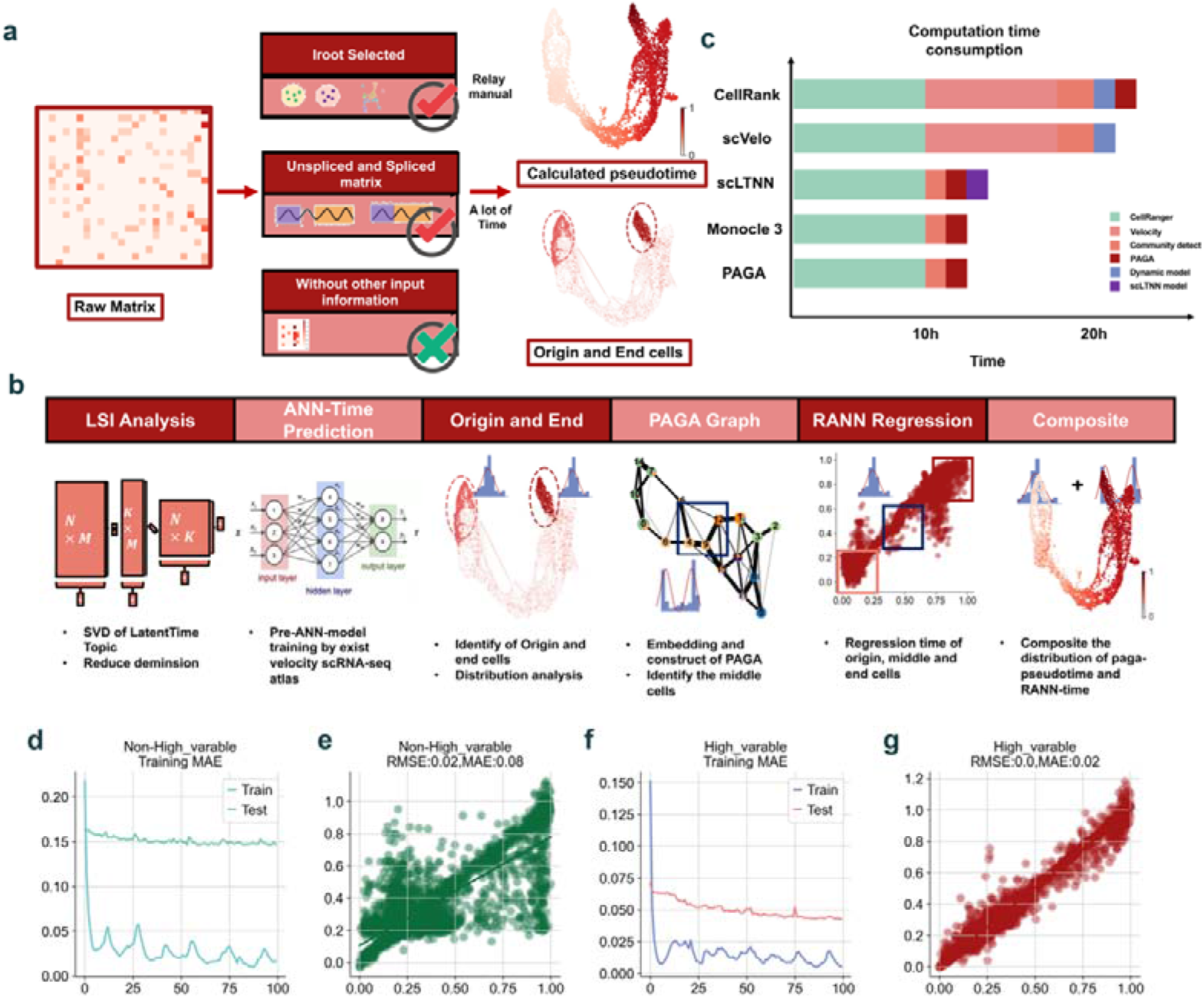
Architecture of the scLTNN framework. (a). Input requirements for calculating pseudotime. (b). scLTNN uses the raw matrix to calculate the origin and end cells by ANN-time prediction model and automatically selects the origin cells as the root of the PAGA graph. The scLTNN then constructed a RANN regression model to predict the intermediate moments using the LSI vectors of cells at origin, middle, and end cells. By compositing the distribution of the RANN-time and PAGA-pseudotime, we calculated the LTNN time to estimate the latent time of the cells. (c) The computation time consumption by different pseudotime calculated methods. (d). The variation of mean absolute error (MAE) during the ANN-time prediction model training by Non-High-variable genes. (e) The Regression analysis of the model predicted results by Non-High-variable genes versus accurate latent time of cells. (f) The variation of mean absolute error (MAE) during the ANN-time prediction model training by High-variable genes. (g) The Regression analysis of the model predicted results by High-variable genes versus true results.

## Results

### LTNN predicts the origin and end cell states automatically with the pre-ANN and CytoTRACE geneset model

The LTNN algorithm aims to model a system’s latent cell time and state. As in the original framework, we model cell latent time using the pre-velocity result by scVelo in different species. For each species, we train an artificial neural network (ANN) model using the latent time of cells and the latent semantic indexing (LSI) of genes. To achieve the best fit, we calculated the LSIs of highly variable and non-highly variable genes separately and used the LSI results for ANN model construction separately. We found that the model fitted by the highly variable genes had better RMSE and MAE (Fig.1d-1g). We also test the different LSI and Batch sizes influenced by the model fitting (sFig.1). We assume that genes in the vast majority of tissues or organs in an organism system have similar distribution characteristics, and thus the ANN model with latent time constructed from single-tissue scRNA-seq can be applied to multiple tissues. To verify our hypothesis, we selected the top 10,000 highly variable genes in 25 human organs and calculated the standard deviation, maximum, median, and mean distribution characteristics of each of these 10,000 genes. We found that the distribution of these 10,000 genes’ standard deviation, median value and mean value were generally consistent in 25 organs, but the distribution of maximum values differed (sFig.2a-2d). To further assess the gene expression features of different organs, we compared the concentration intervals (median divide max value) and standard deviation distributions of the data from different organs (sFig.2e-2f). We have now obtained an ANN model that can evaluate latent cellular time, although the direction of latent time has not yet been determined. We use the expression patterns mentioned in CytoTRACE with genes whose expression patterns correlate with gene counts that may better capture the differentiation state^12^. We use genes whose expression patterns correlate with gene counts to determine the direction of latent time and thus automatically identify cells located in the initial, terminal state of the cell differentiation trajectory. An accurate state prediction method should produce accurate alignment not only in special species but also in different species. Exploiting the true cell state prediction in a true dataset, we further quantified the CD8+ T cell trajectory in human, Pancreatic epithelial, and Ngn3-Venus fusion (NVF) cells during the secondary transition in mouse, 12 time-points spanning 3.3 - 12 hours past fertilization in zebrafish. The LTNN alignment successfully revealed the origin and end cell states in different species. The true annotation of the identified origin cells is initial T cells, and the true annotation of the end cells is depleted T cells in humans (Fig.2a-2c). The true annotation of the identified origin cells is Ductal cells, and the true annotation of the end cells is Beta cells in mice (Fig.2d-2f). The true annotation of the identified origin cells is 04.3 DOME cells, and the true annotation of the end cells is 12.0-6 Somite cells in zebrafish (Fig.2g-2i).

**Figure 2.**
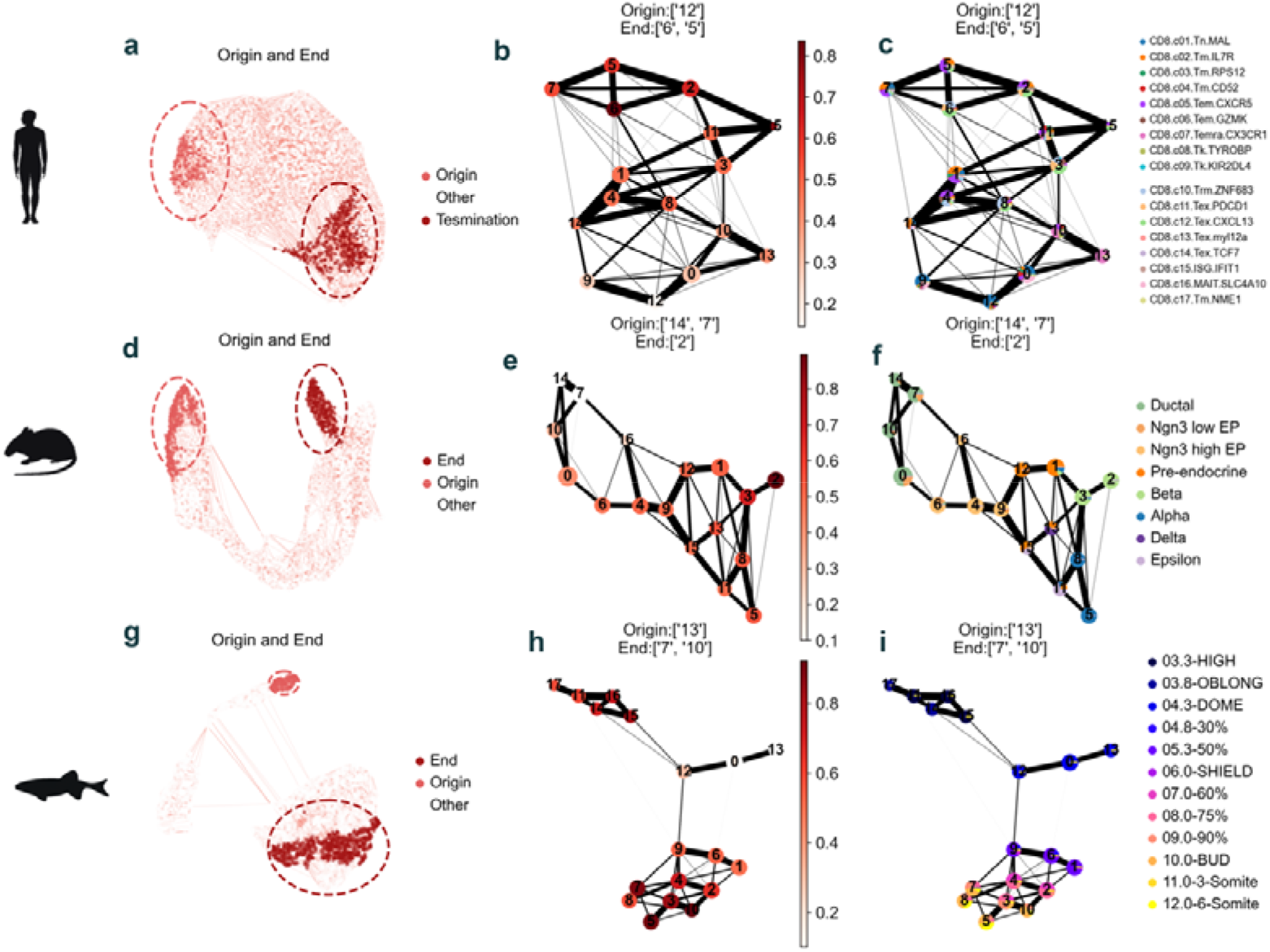
Identify the origin and end cell cross-species. (a). Capture the origin and end cell in human tissue. (b) The score of cellular state by ANN-time prediction model in human tissue. (c). The component of cellular type in a different cellular community in human tissue. (d). Capture the origin and end cell in mouse tissue. (e) The score of cellular state by ANN-time prediction model in mouse tissue. (f). The component of cellular type in a different cellular community in mouse tissue. (g). Capture the origin and end cell in zebrafish tissue. (h) The score of cellular state by ANN-time prediction model in zebrafish tissue. (i). The component of cellular type in a different cellular community in zebrafish tissue.

### Composite regression neural network model based on dweibull distribution to predict the latent time underlying cellular fate

To reconstruct the cell fate trajectory, we calculated a PAGA graph using the cell communities identified by Leiden and calculated the diffusion pseudotime by the origin cell identified by LTNN. We fitted the pseudotime distribution of PAGA and found that it constitutes a dweibull distribution (Fig.3b). In contrast to the three-peaked distribution of latent time of scvelo (Fig.3a), it does not accurately reflect the intermediate state characteristics of the cell fate trajectory. To solve this problem, we selected the origin cells and end cells identified by LTNN and the middle cells identified by PAGA, initialized the time series using the normal distribution for the cells at each of the three-time points, and then constructed an ANN regression neural network to calculate the regression time. We found that the reconstructed regression times also satisfied the normal distribution (Fig.3c). We calculated the probability density function of the normal distribution over Dweibull using the distribution characteristics of the regression time and pseudotime, respectively. We also used the weights of Dweibull for the regression time to enhance the peaks at both ends of the regression time and the weights of the normal distribution for the pseudotime to enhance the peaks in the middle of the pseudotime. We summed the processed regression time and pseudotime to obtain the LTNN time (method). We then evaluated the RMSE and MAE of scVelo’s latent time versus our LTNN’s LTNN time to determine the accuracy of our calculated latent time. We found that the RMSE was 0.01 and the MAE was 0.07 (Fig.3d). The LTNN time is similar to the scVelo’s latent time.

**Figure 3.**
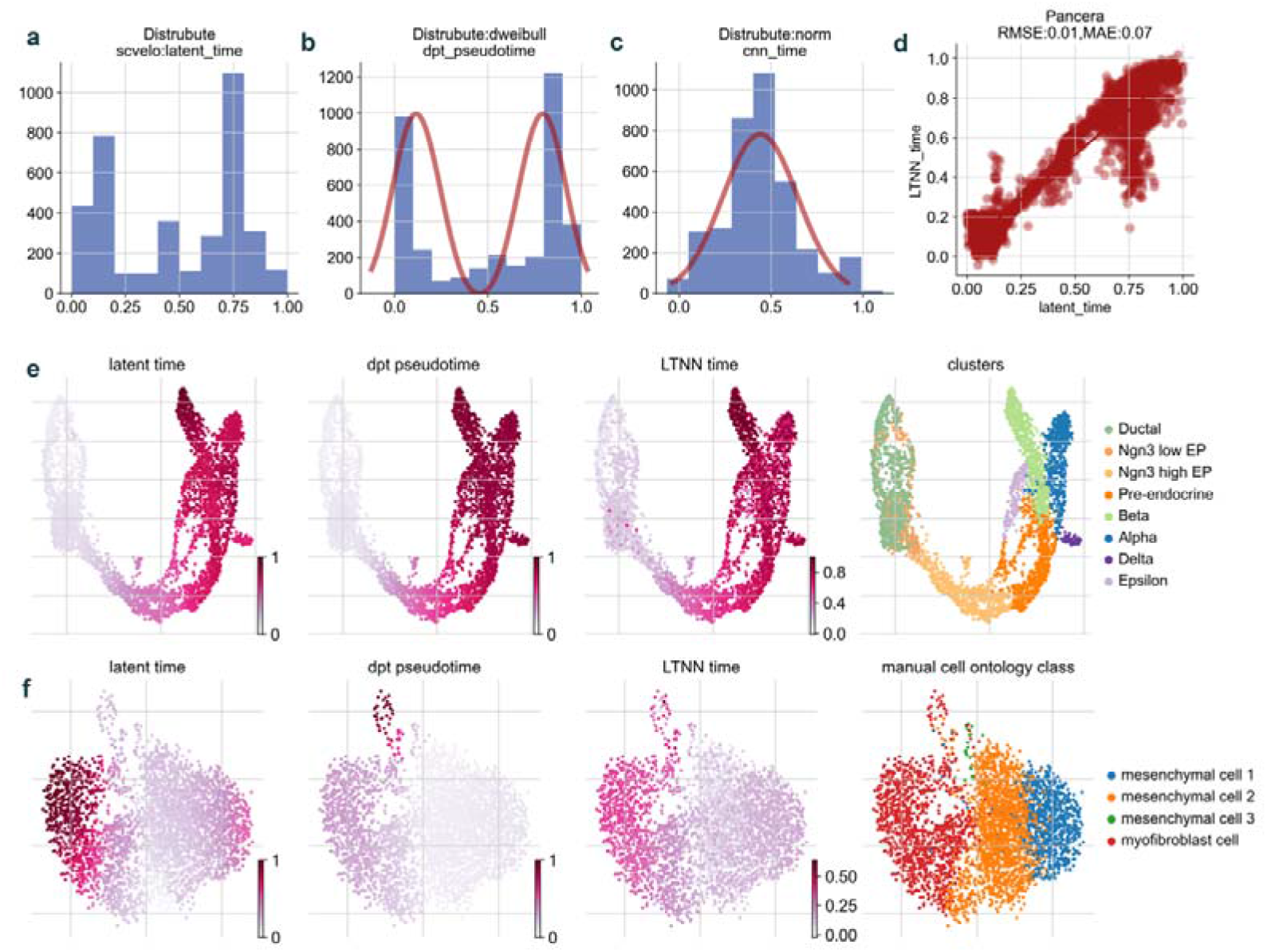
The evaluation of the latent time predicted by scLTNN. The distribution of latent time (calculated by scVelo) (a), dpt_pseudotime (calculated by scanpy) (b), and LSI-NN time (calculated by LSI-NN model) (c) of the pancreas. (d) The Regression analysis of predicted results by the scLTNN model versus latent time results by scVelo. (e) The comparison of latent time, dpt_pesudotime, and LTNN time in different cell types in the pancreas. (f) The comparison of latent time, dpt_pesudotime, and LTNN time in different cell types in bladders.

To evaluate whether LTNN time can overcome the problem of diffusion time and achieve similar results to latent time on a real dataset. We found that LTNN time reflected the Ductal and Beta cell cluster with a similar result of latent time. However, diffusion pseudotime cannot distinguish between Beta and Alpha cells in chronological order in the mouse pancreas dataset (Fig.3e). Similarly, LTNN time distinguish myofibroblast cell well like latent time compared diffusion pseudotime (Fig.3f).

### Identify the dynamical geneset and function in the trajectory of cells cross-species by LTNN

LTNN computes a correlation with LTNN-time for each gene and identifies the dynamical geneset. The geneset allows us to analyze the genes, functions, and pathways involved in cell fate trajectories and thus reveal the key molecular mechanisms behind fate. We have analyzed the molecular mechanisms associated with cell fate in different species. In humans, we analyzed CD8+ T cells, and we found that the expression of ribosome-associated proteins (RPS and RPL families) decreases during the transition from initial T cells to depleted T cells (Fig.4a). While the expression of Cystatin-Like Metastasis-Associated Protein (CST7), CAMP-Responsive Element Modulator (CREM), and C-X-C Motif Chemokine Receptor 4 (CXCR4) proteins was constantly increased. Further functional analysis revealed that membrane protein function, as well as nuclear transcriptional function, were mainly affected during the T cell transformation process (Fig.4d). In mice, we analyzed pancreatic epithelial and Ngn3-Venus fusion (NVF) cells during the secondary transition. We found that the expression of ribosome-associated proteins (RPS and RPL families) also decreases during the transition (Fig.4b). And functional analysis also revealed that membrane protein function, as well as nuclear transcriptional function, were mainly affected (Fig.4e). In zebrafish, we analyzed 12 time-points spanning 3.3 - 12 hours past fertilization and the expression of ribosome-associated proteins (RPS and RPL families) increase during the transition (Fig.4c). The functions are focused on both DNA replication as well as the cell cycle, which is compatible with the development of zebrafish cells (Fig.4f).

**Figure 4.**
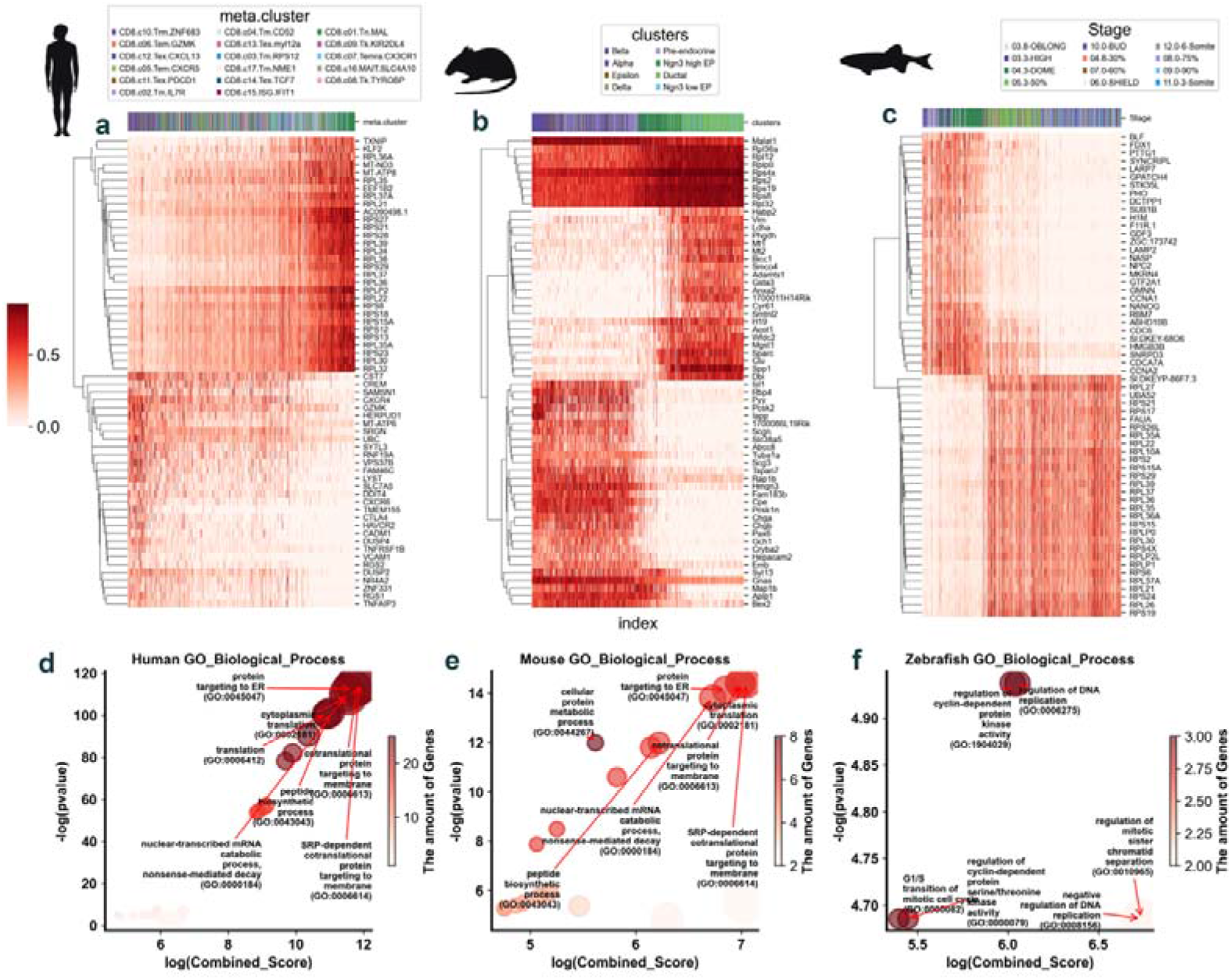
The expression and function of key genes in the trajectory of cells cross-species. (a) The expression of top 30 key genes in the trajectory of human tissue. (b) The expression of top 30 key genes in the trajectory of mouse tissue. (c) The expression of top 30 key genes in the trajectory of zebrafish tissue. (The value of expression was performed normalization by min-max) The Gene ontology biological process of top 30 key genes in the trajectory of human tissue(d) mouse tissue (e) zebrafish tissue (f). (p-value<0.05, the shade of color display the number of genes in different GO term)

## Discussion

scLTNN enables the latent time of cell estimation, origin and end cell identification, and cell fate trajectory computation without additional information. On the one hand, it reduces a large amount of consumption of computer resources and waiting time. On the other hand, it enables researchers to make cell trajectory inferences without a very sophisticated biological background. Novelty, scLTNN enables the development of cell trajectory inference analysis in the same type of cells. It is difficult for researchers to identify the start node of the trajectory without RNA velocity in regeneration, reprogramming, and cancer.

We note that the current framework is cross-species supported. scLTNN accurately identifies origin andend cell states from scRNA-seq data by combining a priori latent time predictions and genes whose expression patterns correlate with gene counts. We tested the ability of scLTNN on real data sets derived from different species separately. In addition, we successfully constructed a proposed temporal trajectory inference algorithm comparable to scVelo’s latent time by combining Dweibull, a spectral distribution feature constructed by PAGA, and Norm, a spectral distribution feature of origin, middle and end cell states.

Beyond the cell lineage tracing and kinetic process analysis of relevant genes, dynamic activation of pathways as well as functions is also important. By combining scLTNN with the enrichment method, activated pathways and functions can be inferred in biological processes, without any differential expression analysis, providing another idea for studying cell fate dynamics.

We believe that scLTNN, as a simple and flexible framework without any additional information, creates an unprecedented opportunity for the inference of cell fate trajectory and cell state. The whole package of scLTNN, along with tutorials and demo cases, is available online at https://github.com/Starlitnightly/scLTNN for the community.

## Materials and Methods

### Preparing the scRNA-seq data for LSI vector calculation

The raw data set of pancreatic epithelial and Ngn3-Venus fusion (NVF) cells during secondary transition^13^ is available in the National Center for Biotechnology Information’s Gene Expression Omnibus repository under accession number GSE132188. We included samples from the last experimental time point: E15.5. The raw data set of mesenchymal to myofibroblast in the bladder^14^ has been deposited under accession number GSE201333. We included samples from the CD8+ T cell in cancer^15^. The raw data set of has been deposited under accession number GSE156728. We included samples from the sample of TSP1. The count matrices are size normalized to the median of total molecules across cells. The top 10,000 highly variable genes are selected by the Seurat_v3 method in the Python library scanpy^16^. The non-highly variable genes were obtained by taking intersections of genes that recur in 10% of the cells in 25 human organs.

### Latent semantic indexing neuron network framework

The latent semantic indexing neuron network (LSI-NN) was constructed by two structures. We assume that there are *m* cells, each cell with a *n* genes to be indexed. *A*_*m*×*n*_ denotes the matrix prepared for LSI analysis^17^. We use SVD decomposition to decompose the matrix *A*_*m*×*n*_ into the following three matrices:

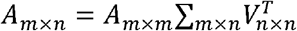

We then assume there are *k* topics and perform rank reduction on the SVD decomposition:

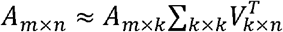

In the scRNA-seq data, *t*_*m*_ denotes the timing sequence of the cells, which is also present as a latent semantic. We can use *A*_*m*×*k*_ and *t*_*m*_ to construct a regression model. LSI computed using the scglue library in Python^18^.

We assume K (K=5) layers in the artificial neural network^19^. *Net*^(*K*)^ denotes all nodes in the K layer. *Y*^(*K*)^ denotes each layer’s output. To learn the regression model, we minimize the following objective function:

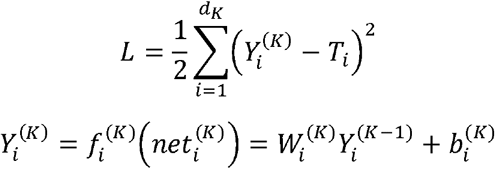

Where 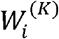 indicates the weight of node *i* in K layer, 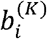 is the bias of node *i* in K layer, 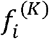 denotes the activate function in neuron, which is Relu in the hidden layer, and Linear in the output layer. Minimization is obtained via gradient-based optimization using the Keras library in Python^20^. To evaluate the consistency of regression result 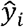 and true result *y*_*i*_, we use mean absolute error(MAE).

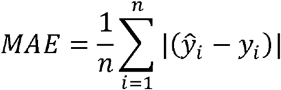

### Latent time neuron network framework

We assume the diffusion pseudotime follows the double weibull distribution (*X*~*dweibull*(*k*, *μ*, *σ*))^21^, and the LSI-NN time follows the norm distribution (*X*~*N*(*μ*, *σ*^2^))^22^. The latent time neuron network (LTNN) was constructed by two distributions:

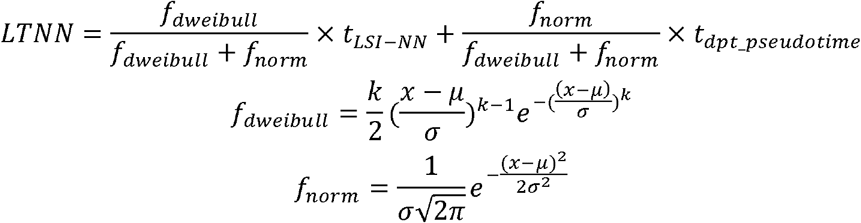

To estimate the k, *μ*, and *σ* of dweibull and norm distribution, maximum likelihood estimation using the scipy library in Python^23^.

### Comparison of the highly variable gene expression feature in multiple-organ of humans

We used the multiple-organ scRNA-seq atlas of humans to perform the gene expression feature test. The data has been deposited at figshare. A total of 25 organ data were used in this study^24^. After selecting the top 10,000 highly variable genes by Seurat_v3^16^, we then calculated the max, mean, and median expression of genes in each cell. Hist plots were used to visualize the distribution of max, mean and median expression. Standard deviation was used to evaluate the feature of distribution of max, mean and median expression.

### Trajectory inference and PAGA graph construction for scRNA-seq

We used Leiden to cluster the cell, capture the underlying topology, and used PAGA to compute the unified pseudotime by averaging the single-cell level diffusion pseudotime computed by DPT^10^. We manually specificated the terminal cell fates and clusters by existing research to compare gene expression trends across lineages.

### Identify the dynamical geneset and function

We evaluated the correlation using Pearson’s coefficient for LTNN time for all cells and the expression of each gene in all cells to find the key set of kinetic process genes. And the gene ontology (GO_Biological_Process_2018)^25^ of the gene sets was evaluated using the Python package pyGSEA(v.0.10.4, https://pypi.org/project/gseapy/).

## Supporting information

sFig.1, sFig.2

## Data availability

All datasets used in this study are already published and were obtained from public data repositories. The raw data set of pancreatic epithelial and Ngn3-Venus fusion (NVF) cells during secondary transition^13^ is available in the National Center for Biotechnology Information’s Gene Expression Omnibus repository under accession number GSE132188. The raw data set of CD8+ T cell^26^ has been deposited under accession number GSE156728. We included samples from the last experimental time point: E15.5. The raw data set of mesenchymal to myofibroblast in the bladder has been deposited under accession number GSE201333. We included samples from the sample of TSP1.

## Code availability

The scLTNN framework was implemented in the scltnn Python package, which is available at https://github.com/Starlitnightly/scltnn. For reproducibility, the scripts for all training and evaluation were introduced by the Jupyter notebook, which is also available in the above repository.

## Acknowledgments

This work was supported by the Hebei Provincial Department of Science and Technology (No.19942410G). Cell Therapy Laboratory, the First Hospital of Hebei Medical University, Shijiazhuang, Hebei 050031, China; Department of Immunology, Basic Medical College, Hebei Medical University, Shijiazhuang, Hebei 050017, China. Electronic address.

This work was supported by the Student Research Training Program of University of Science and Technology Beijing. This work was also supported by the Doctoral Research Fund of the University of Science and Technology Beijing (No. 06198268).

## Author contributions

Z.Z. Conceived, designed, and developed the method, implemented scLTNN, analyzed the data and wrote the Methods and Result section. C.X. interpreted the relevance of the method for inferring development trajectories in the zebrafish data. L.H. and R.S. contributed to the implementation and reproduction test. Y.X. and H.D. supervised the project and contributed to the conception of the project. Z.Z, L.H. and C.X. wrote the manuscript with contributions from the co-authors. All authors read and approved the final manuscript.

