## Supplementary material for "Identify the origin and end cells and infer the trajectory of cellular fate automatically": sFig.1, sFig.2

### Supplement materials

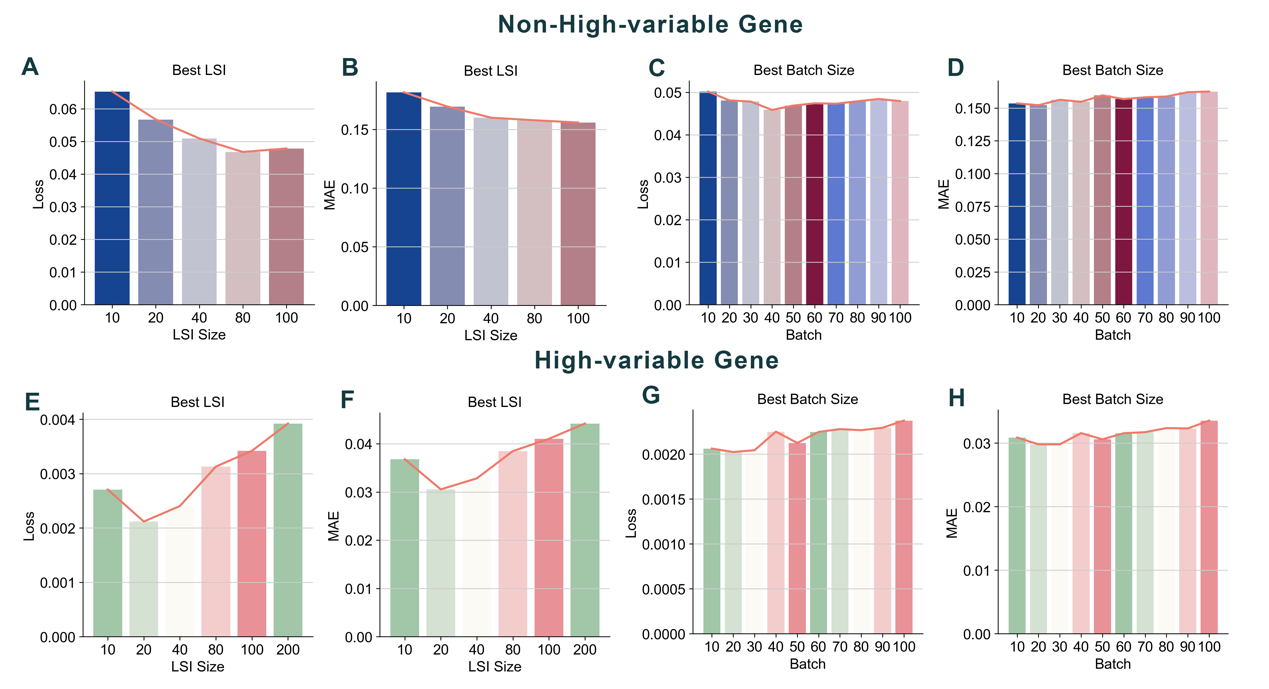

sFig 1 | The Selection of the best superparameters during LSI-CNN model training. Non-High-variable Gene: (A) The evaluation of best LSI components by loss. (B) The evaluation of best LSI components by MAE. (C) The evaluation of best batch size by loss. (D) The evaluation of best batch size by MAE. High-variable-Gene: (E) The evaluation of best LSI components by loss. (F) The evaluation of best LSI components by MAE. (G) The evaluation of best batch size by loss. (H) The evaluation of best batch size by MAE.

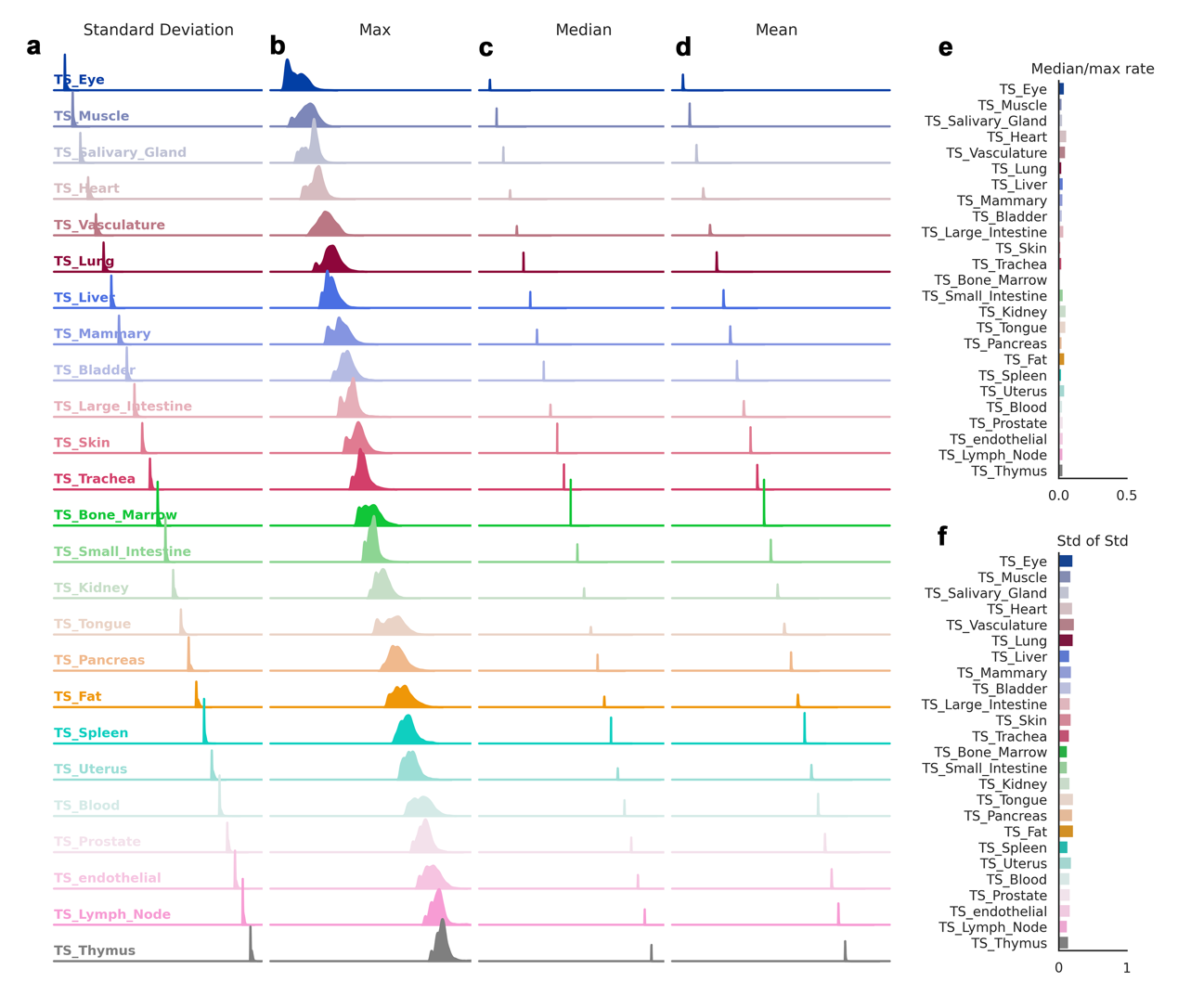

**sFig.2 | Expression distribution of multiple-organ scRNA-seq atlas of human.** The standard deviation (a), Max value (b), Median value (c) and Mean value (d) of the top 10,000 high-variable genes in 25 human organs. (e). The rate of median value divide by max value of 10,000 high-variable genes in 25 human organs. (f). The standard deviation of standard deviation in 10,000 high-variable genes of 25 human organs.
